# Factors influencing the detergent-free membrane protein isolation using synthetic nanodisc-forming polymers

**DOI:** 10.1101/2023.05.12.540572

**Authors:** Bankala Krishnarjuna, Gaurav Sharma, Thirupathi Ravula, Ayyalusamy Ramamoorthy

**Author notes:** **Corresponding Author:** Ayyalusamy Ramamoorthy. equal contribution (Bankala Krishnarjuna, Gaurav Sharma).

## Abstract

The detergent-free isolation of membrane proteins using synthetic polymers is becoming the desired approach for functional and structural studies of membrane proteins. Since the expression levels for many membrane proteins are low and a high yield of functionalized reconstituted membrane proteins is essential for *in vitro* studies, it is crucial to optimize the experimental conditions for a given polymer to effectively solubilize target membranes/proteins. The factors that affect membrane solubilization and subsequently the isolation of a target membrane protein include polymer concentration, polymer charge, temperature, pH, and concentration of divalent metal ions. Therefore, it is important to have knowledge about the efficacy of different types of polymers in solubilizing cell membranes. In this study, we evaluate the efficacy of inulin-based non-ionic polymers in solubilizing *E. coli* membranes enriched with rat flavin mononucleotide binding-domain (FBD) of cytochrome-P450-reductase (CPR) and rabbit cytochrome-b5 (Cyt-b5) under various solubilization conditions. Our results show that a 1:1 (w/w) membrane:polymer ratio, low temperature, high pH and sub-millimolar concentration of metal ions favor the solubilization of *E. coli* membrane enriched with FBD or Cyt-b5. Conversely, the presence of excess divalent metal ions affected the final protein levels in the polymer-solubilized samples. We believe that the results from this study provides knowledge to assess and plan the use of non-ionic polymers in membrane protein studies.

## 1. Introduction

There is considerable interest in the functional reconstitution and characterization of membrane proteins in a native lipid membrane environment without the use of any detergents [1-7]. Recent studies have successfully developed approaches for the detergent-free isolation of membrane proteins and lipids from the cell membrane, and to form nanodiscs for *in vitro* studies [8-11]. Several synthetic, lipid-solubilizing polymers such as ionic styrene–maleic acid (SMA; and its derivatives, including SMA-QA and SMA-EA), diisobutylene maleic acid (DIBMA; and their modified versions), polymethacrylate, zwitterionic SMA and non-ionic inulin-based polymers have been successfully used to isolate membrane proteins without the use of any detergents [12-22]. Moreover, several high-resolution structures have been determined for membrane proteins that are directly isolated using detergent-free membrane protein isolation strategies [23-25]. Due to high-charge density, ionic polymers have limitations in studying membrane proteins under certain solubilization conditions [17, 26-28]. The non-ionic polymers are developed to overcome some of the limitations posed by ionic polymers to study membrane proteins containing large soluble domains and their complexes [10, 17, 26, 29].

Since many membrane proteins do not express in high yield and a significantly large amount of functionalized reconstituted membrane proteins is desired for *in vitro* studies, it is crucial to optimize the conditions for a given polymer to effectively solubilize the target cell membranes [28, 30-33]. Several factors, including polymer length, polymer type, sample temperature, concentration of metal ions and pH of buffer affect the solubilization of cell membranes and, subsequently, the target protein yields [28, 31, 34, 35]. The aqueous solubility of a polymer itself is another key factor that affects polymer’s efficacy in solubilizing cell membranes [31]. SMA has two acidic groups per maleic acid unit with different pKa values (pK_1_∼5.5, pK_2_∼8.6) [31]. Hence, at acidic pH (pH < 5), polymer self-assemble into large aggregates due to increasing protonation of maleic acid groups, whereas in alkaline pH, the polymer is highly hydrophilic due to high-charge density that makes it less efficient in solubilizing cell membranes [31]. SMA 2:1 is a more promising polymer to work at a wide pH range compared to other versions of SMA polymers. In addition, due to high-charge density, SMA polymers are susceptible to divalent metal ions; they precipitate in the presence of even low mM concentrations of Mg^2+^ (∼10 mM) and Ca^2+^ (∼5 mM) ions [28, 36]. DIBMA has been shown to work better at these concentrations of divalent ions [36]. Short-length SMA or DIBMA polymers are more efficient than high-molecular-weight polymers [35]. All the mentioned conditions and their effects can vary from one polymer to another. Therefore, it is important to evaluate how the different sample conditions affect the membrane solubilization properties of non-ionic polymers.

In this study, we used pentyl-inulin (inulin functionalized with a hydrophobic pentyl group) to systematically evaluate its efficacy in solubilizing *E. coli* cell membranes enriched with a single transmembrane domain-containing rat flavin mononucleotide binding-domain (FBD) of cytochrome P450 reductase (CPR) or rabbit cytochrome b5 (Cyt-b5) proteins (see the Supplementary Information for protein sequences). CPR and Cyt-b5 are redox partners of cytochrome-P450 and play important roles in the drug metabolism by cytochrome-P450 enzymes [37-39]. The solubilization experiments were performed in the presence of various buffer environments, including temperature, divalent metal ions, pH, polymer concentration and polymer functionalized with different types of hydrophobic groups (**Fig. 1**). The extent of solubilization was determined by measuring protein concentration using absorbance spectroscopy or by quantifying protein band intensities on sodium dodecyl-sulfate polyacrylamide gel electrophoresis (SDS-PAGE) gel to test the efficacy of the polymer.

**Figure 1.**
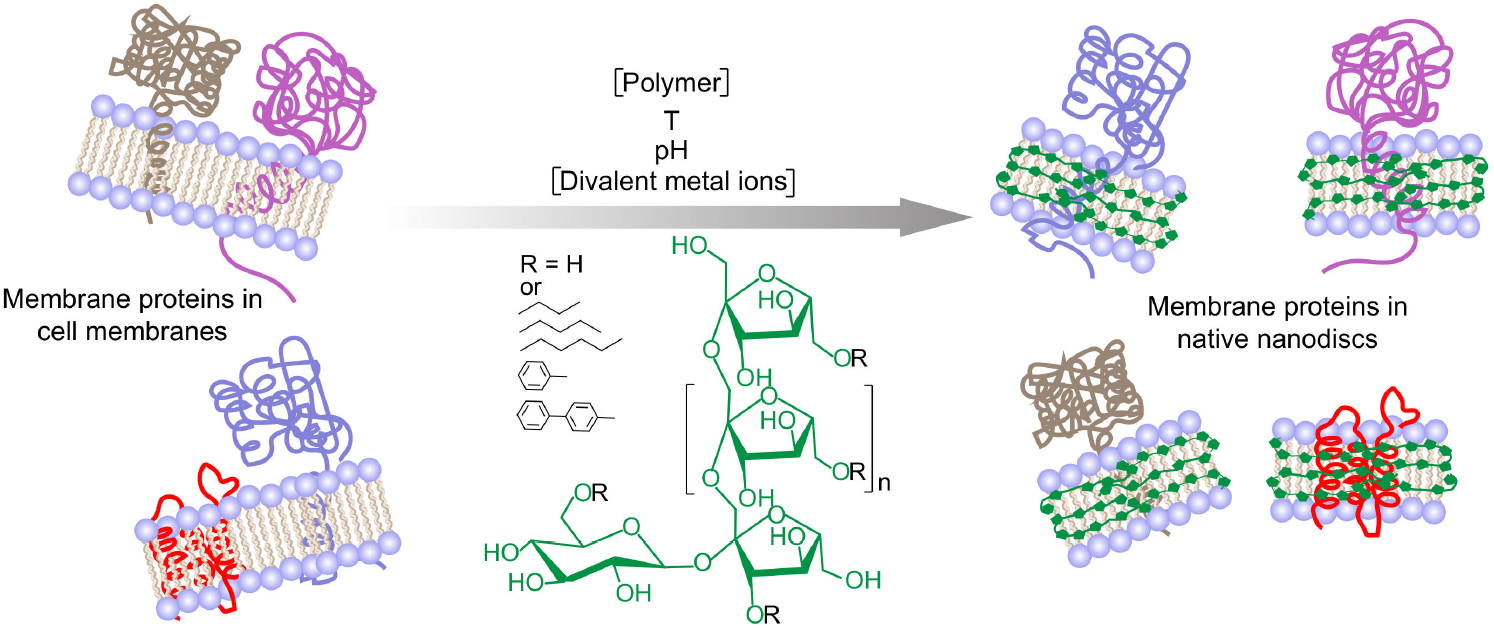
Schematic of the solubilization of cell membranes using a synthetic inulin-based polymer under different solubilizations conditions, including the concentrations of the polymer and divalent metal ions (such as Ca^2+^ and Mg^2+^), temperature and pH. Reconstitution of membrane-bound proteins and other membrane components directly isolated from the cell membrane in polymer nanodiscs shown on the right. Five different synthetic non-ionic inulin-based polymers that differ in the functionalization of the hydrophobic group (R: butyl, pentyl, hexyl, phenyl, biphenyl) were used in this study. The polymer structure/polymer-belt are shown in green. Only the membrane protein-containing nanodiscs are shown, and other insoluble membrane components are omitted for clarity (right).

## 2. Materials and Methods

### 2.1. FBD and cytochrome-b5 expression in E. coli cells

The synthetic genes encoding Cyt-b5 and FBD were purchased from GenScript and subcloned into pET28a(+) vector (restriction sites; NcoI/XhoI). The *E. coli* (C41) cells for protein expression were purchased from Lucigen Corporation (Wisconsin). These proteins were overexpressed using the published protocol [10, 17, 40]. Kanamycin (Thermo Fisher Scientific, Waltham, MA, USA) was used as a selection marker. When the culture OD_600_was ∼0.8 (FBD) or 0.5 (Cyt-b5), the protein overexpression was induced by adding 0.5 mM (Cyt-b5) or 0.4 mM (FBD) isopropyl ß-D-1-thiogalactopyranoside (IPTG) (Sigma-Aldrich). Cyt-b5 was overexpressed at 37 °C for ∼18 h and FBD was overexpressed at 30 °C for ∼16 hours. Cells were harvested by centrifugation at 6000 rpm for 10 min.

### 2.2 Cell lysis and membrane preparation

Cell pellets were resuspended in 50 mM Tris buffer (pH 7.4) containing 100 mM NaCl (Fisher Scientific) and a cOmplete™ protease inhibitor cocktail tablet (Sigma-Aldrich)). Cells were lysed by incubating them with lysozyme (Sigma-Aldrich) (5 mg/g wet cell paste), DNase (Sigma-Aldrich) (0.1 mg/g wet cell paste), and 2 mM MgCl_2_ (Sigma-Aldrich), followed by 10 cycles (each 10 seconds pulse) of sonication with 1 min interval between the successive cycles. First, the soluble components of lysate were removed by centrifugation at 18,000 rpm for 60 min. Then the membranes were washed by resuspending in 10 mM Tris (pH 7.4) buffer containing 100 mM NaCl and protease inhibitors. Finally, upon centrifugation, the washed membranes were used to make the membrane stock solution at 250 mg/mL concentration. All the steps were performed on ice.

### 2.3. Polymer synthesis

Inulin-polymers functionalized with different hydrophobic groups were synthesized by using the published protocols [34]. Briefly, 1 g of inulin was added to 30 mL of dimethylacetamide (DMAc) and heated to 50 °C with stirring until all inulin was solubilized (∼1 hr). The reaction was then allowed to cool to room temperature and sodium hydride (3×0.5 eq, 442 g, 60% NaH in mineral oil) was added slowly in 3 portions over the course of 1 hr. The resulting mixture was heated to 50 °C with stirring for an additional hour. The reaction was then allowed to cool, followed by the addition of 1.06 mL of n-bromide (n=butyl/pentyl/hexyl/benzyl/di-benzyl). The reaction was then allowed to stir at room temperature. The degree of functionalization was determined by taking 100 μL of the reaction solution, precipitating with ethyl ether and ethanol (50% v/v), followed by centrifugation (3000 rpm, 5 min) and separation of supernatant from pellet. The pellet was subsequently washed with pure ethyl ether 3 times. It was then allowed to dry under nitrogen gas and subsequently dried under high vacuum. The resulting powder was dissolved in ^2^H_2_O and characterized by ^1^H NMR (without water suppression). The reaction was monitored for 2-3 days until the desired degree of functionalization (∼33 %) was achieved. Once achieved, the reaction was stopped by addition of cold ethanol and ethyl ether (50% v/v). The precipitate was separated through centrifugation and washed three times using ethyl ether, as well as washed 3 times using ethanol. The precipitate was then dried under vacuum. The resulting white powder was then dissolved in water and dialysis was done using 1 kDa MWCO dialysis tubing to remove the impurities. The sample was then lyophilized to obtain a white powder. The polymer was characterized using ^1^H NMR (without water suppression).

### 2.4. Membrane solubilization using inulin based polymers

The membrane solubilization experiments were performed using *E. coli* membranes at 25 mg/mL final concentration. For the temperature-dependent experiments, three samples containing membranes and polymer at a 1:1 (w/w) ratio were prepared on ice. The samples were then incubated overnight at three different temperatures: 4, 25, and 37 °C with a gentle mixing. To test the effect of the synthetic polymer, four different samples with 0.2:1, 0.4:1, 0.7:1, and 1:1 (w/w) polymer-to-membrane ratios were prepared and incubated at 4 °C overnight with a gentle mixing. For pH-dependent experiments, the membranes were resuspended in pH 7, 8, and 9 buffers containing 50 mM Tris and 100 mM NaCl, and the solubilization was performed at 4 °C overnight. To see the effect of divalent metal ions, the samples contained 5 different concentrations of MgCl_2_and CaCl_2_were prepared. After the solubilization of cells using the polymer, the insoluble material was removed by centrifugation at 10000 rpm for 45 min, and the soluble fraction was used for further analysis.

### 2.5. SDS-PAGE analysis of polymer-solubilized cell membranes

The soluble fractions from polymer-solubilized samples were analyzed by SDS-PAGE. The samples were mixed with 2x dye at a 1:1 v/v ratio and heat treated at 90 °C for 4-5 minutes before loading onto the gel. The pre-cast gradient gels (4-20 %) were purchased from GenScript (Piscataway, NJ). Electrophoresis was performed using a Bio-Rad SDS-PAGE gel analysis casting system (Bio-Rad Laboratories, Inc.; Hercules, CA, USA) operated at 150 volts. The gels were stained for 1 h in a staining solution (40 % ethanol, 10 % acetic acid and 0.25 % Coomassie Brilliant Blue R-250 (ThermoScientific)). The gels were destained using the same solvent mixture without Coomassie Brilliant Blue R-250, and the gels were stored in Milli-Q water before imaging.

### 2.6. Quantification of membrane proteins in the polymer-solubilized samples

The destained SDS-PAGE gels were scanned using the Image Lab^™^ touch software (version 2.4.0.03) in the ChemiDocTM MP Imaging System (Bio-Rad). The FBD, Cyt-b5 and other protein band intensities were determined using the volume contour tool Image Lab (Bio-Rad; Hercules, CA, USA) or ImageJ software. For all the data reported in this study, all the protein bands from every independent sample were analyzed in duplicate pairs of solubilized fractions, with each pair being loaded on the same gel.

### 2.7. Absorbance spectra of cytochrome-b5 in the polymer-solubilized samples

The Cyt-b5 samples were analyzed by recording absorbance spectra between 380-600 nm using UV/vis spectrophotometer (DeNovix, DS-11+). The data were collected on freshly prepared solubilization samples. The samples without the polymer were used to record reference spectra. The spectra were recorded in the presence and absence of the reducing agent sodium dithionite (Sigma-Aldrich). The protein concentration was measured using the Beer-Lambert formula: A=εcl (A= absorbance, ε= molar extinction coefficient, c=protein concentration and l= path length (or the dimension of the sample cell). A molar extinction coefficient, ε, of 185 mM^−1^ cm^−1^ for the absorbance change (i.e. the intensity at 423 minus the intensity at 409 nm) was used to calculate the amount of Cyt-b5 present in the unpurified/polymer-solubilized samples [40].

## 3. Results and Discussion

### 3.1. E. coli membrane solubilization improved with increasing pentyl-inulin concentration

Membrane solubilization was tested using four different concentrations of pentyl-inulin by keeping the amount of membranes constant: 1:0.2, 1;0.4, 1:0.7, and 1:1 w/w membrane:polymer ratios. Since Cyt-b5 show characteristic absorbance spectral properties in the visible region [17, 40], we used absorbance spectra in addition to protein densitometric analysis to estimate the amount of Cyt-b5 present in the polymer-solubilized samples. The absorbance spectra were recorded under oxidized and reduced conditions. The intensity of the new peak appearing at 423 nm under the reduced condition increased with the increasing pentyl-inulin concentration (**Fig. 2A**). Cyt-b5 concentration was measured using a difference extinction coefficient of 185 mM ^−1^ cm^−1^ for the absorbance change at 423-409 nm. Cyt-b5 concentration substantially increased when the membrane:polymer ratio was increased from 1:0.2 to 1:0.4 (**Figs. 2B, S1**). The yield of protein isolation further increased by 10% upon changing the membrane:polymer ratio (w/w) from 1:0.4 to 1:1, indicating the membrane solubilization was influenced by the amount of polymer in the sample and the maximum solubilization was observed at a 1:1 w/w membrane:polymer ratio. In addition, the amount of Cyt-b5 was also quantified from SDS-PAGE (**Figs. 2C, S1**). The sample with the membrane:polymer ratio of 1:0.2 showed minimal protein band intensity and when the membrane:polymer ratio (w/w) used at 1:0.4, the protein band intensity substantially increased by 20% (**Fig. 2C**). The observed band intensity was further increased by ∼10% upon changing the membrane:polymer ratio (w/w) to 1:0.7 (**Fig. 2C**). There was no significant difference between the samples with 1:0.7 and 1:1 membrane:polymer ratios (w/w) (**Fig. 2C**). These observations are complementary to the results obtained from absorbance spectra. The membrane sample excluding the polymer did not show any characteristic absorbance peak for Cyt-b5. In the case of FBD, the same trend of solubilization was observed; i.e., the observed FBD band intensity substantially increased when the membrane:polymer ratio (w/w) was increased from 1:0.2 to 1:0.4 or 1:0.7 and there was no significant change for 1:0.7 or 1:1 membrane:polymer ratios (**Figs. 2D, S1**). Since the amounts of Cyt-b5 quantified using absorbance spectra and SDS-PAGE were not substantially different, we used one of these experiments to see the effect of pH, temperature and metal ions on membrane solubilization and protein yields. The optimal concentration of polymer that is required for membrane solubilization might vary from one protein to another as the annular lipid composition can differ from protein to protein or membrane to membrane. In addition, the impurities from chemical synthesis can also affect the overall solubility and the membrane-solubilizing efficacy of a given polymer.

**Figure 2.**
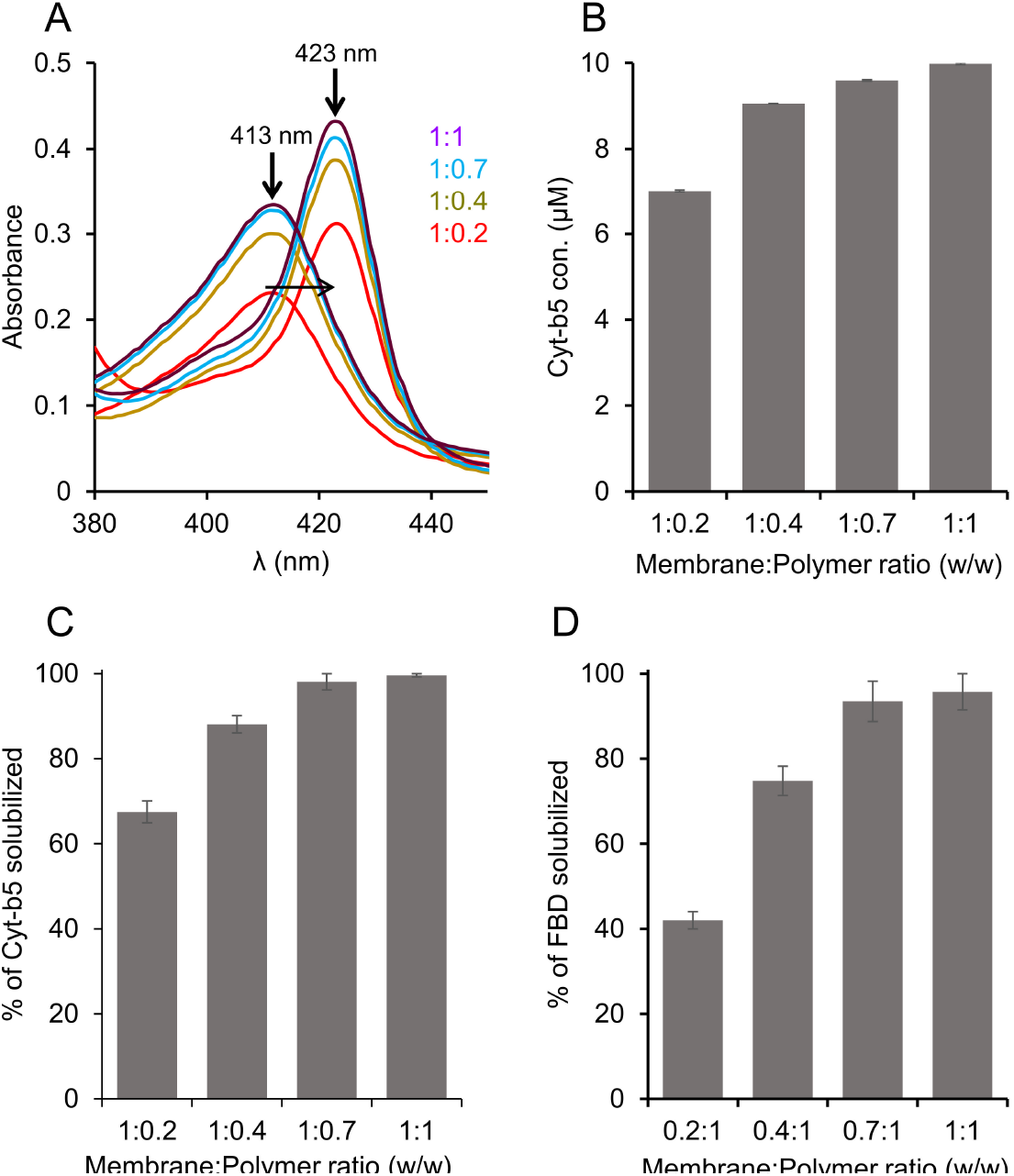
Solubilization of Cyt-b5 or FBD enriched *E. coli* membranes in the presence of different concentrations of pentyl-inulin. The quantity of membranes was kept constant (250 mg/mL), and the amount of polymer added was varied to obtain four different membrane:polymer w/w ratios, 0.2:1, 0.4:1, 0.7:1, and 1:1. The membranes were solubilized at 4 °C overnight. (**A**) Absorbance spectra measured to determine the protein concentration in the polymer solubilized membrane samples. The horizontal arrow indicates the disappearance of peak at 413 nm (oxidized form) and the appearance of new peak at 423 nm (sodium dithionite-reduced form). The increase in the absorbance peak intensity is shown with an upside arrow, and the membrane:polymer ratios (w/w) are indicated. The data shown are from one of the duplicates of solubilization experiments. (**B-D**) Bar plots depicting (B) the concentration of Cyt-b5 determined from (A), the percentage of (C) Cyt-b5 and (D) FBD protein bands (from SDS-PAGE gels) quantified by ImageJ.

### 3.2. Temperature affects pentyl-inulin’s solubilization of E. coli membranes in a protein-dependent manner

The *E. coli* cell membranes enriched with FBD or Cyt-b5 were solubilized using the pentyl-inulin polymer at three different temperatures 4, 25, and 37 °C. The solubilized FBD and the total membrane protein samples were analyzed by SDS-PAGE, while the amount of Cyt-b5 solubilized by the polymer was quantified by absorbance spectra (**Fig. S2**). The protein band intensities for the FBD samples solubilized at 4 °C and 25 °C were similar. But, the FBD protein band intensity was decreased by ∼20 % upon increasing the temperature to 37 °C. Thus, the efficacy of pentyl-inulin solubilizing FBD-rich membranes was effective at lower temperatures than at higher temperatures (**Fig. 3A**). In the case of Cyt-b5, the absorbance spectra showed an increase in the intensity of 409 and 423 peaks at 25 °C than at 4 °C and 37 °C (**Fig. 3B**). As a result, the protein concentration determined from the difference spectra was higher when the membranes were solubilized at 25 °C than at 4 °C or 37 °C (**Fig. 3C**). A similar trend was observed when all the membrane proteins from *E. coli* membrane were quantified (**Fig. 3D**). Overall, these results suggest that FBD/Cyt-b5-rich membranes were better solubilized by the polymer at low temperatures (4 °C/25 °C) than at 37 °C. The difference in the extent of solubilization observed for FBD and Cyt-b5-rich membranes might be due to the difference in the lipid composition surrounding the transmembrane domains of these proteins. For example, the lipid domains rich in non-polar lipids may not be solubilized as efficiently as polar lipids by polymers [41]. Furthermore, the type of other membrane proteins surrounding the target membrane protein in the cell membrane can vary; this difference can also affect the solubilization of the target membrane/protein. Additionally, higher temperatures may enhance the solubility of some of the membrane proteins, but such conditions may become more susceptible to membrane-associated proteases at higher temperatures due to increased dynamics of molecular components present in the cell membrane. Therefore, the membrane-solubilization and the final protein yields can be protein dependent.

**Figure 3.**
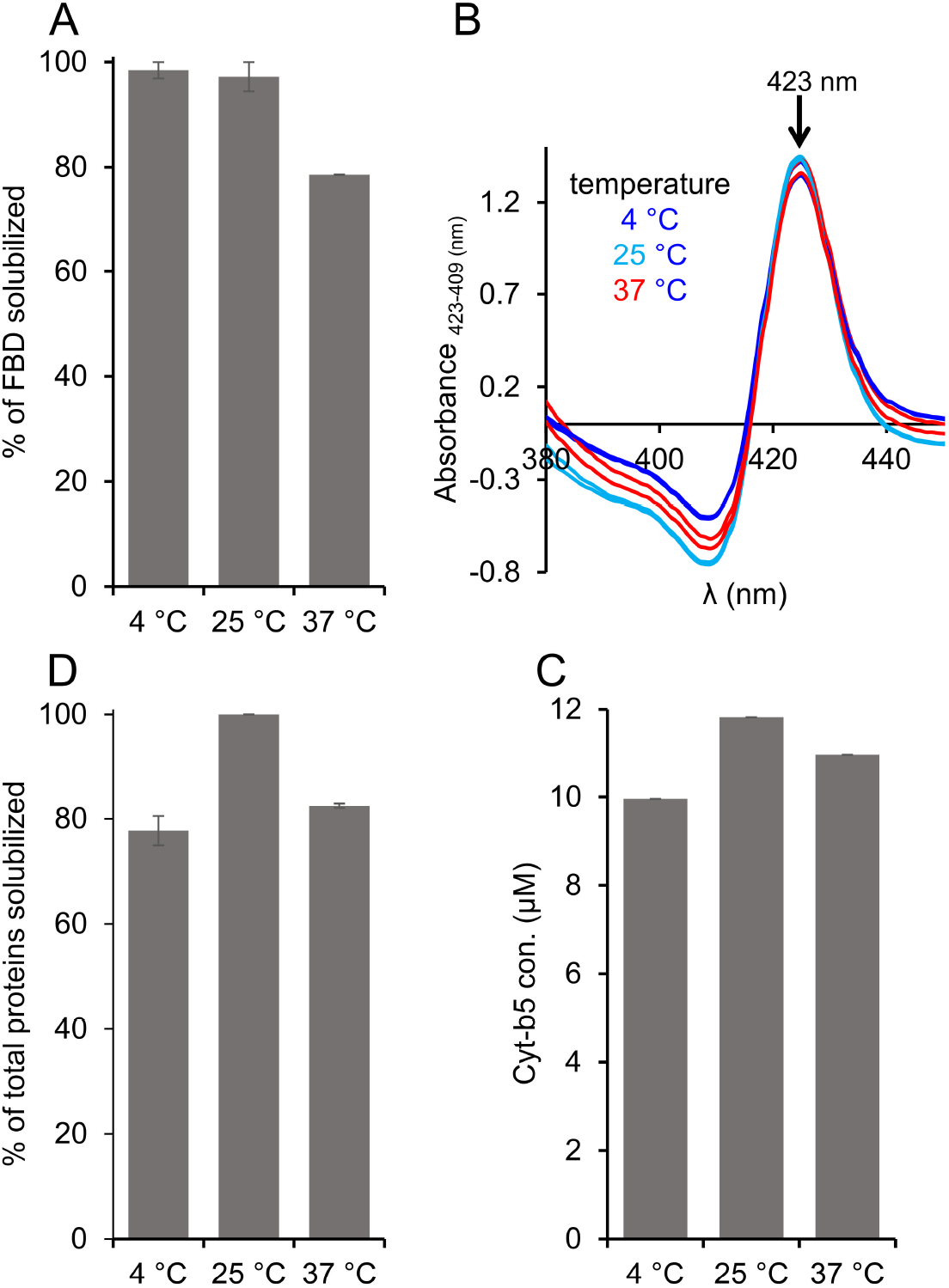
Variable-temperature solubilization of FBD and Cyt-b5-enriched *E. coli* membranes. (**A**) FBD, (**B, C**) Cyt-b5 and (**D**) total membrane proteins analyzed from SDS-PAGE and absorbance spectra. The absorbance difference spectra were obtained by subtracting the absorbance spectra of oxidized Cyt-b5 from that of reduced Cyt-b5 (B). Upside arrow indicates an increase in the absorbance observed at 423 nm. The samples were prepared using 1:1 [w/w] membrane:polymer ratio (25 mg/mL each). All the samples were prepared on ice before incubating them at different temperatures 4, 25, and 37 °C. The samples were analyzed by SDS-PAGE and absorbance spectra. Image Lab software was used to quantify the membrane protein band intensities. The plots were generated from two independent experiments, and error bars indicate the difference in the extent of solubilization measured from them. The data were normalized to the sample that showed the maximum solubilization at each solubilization experiment.

### 3.3. pH affects the pentyl-inulin’s solubilization of E. coli membranes

Buffer pH is one of the key factors that affect the efficacy of polymers in solubilizing the cell membranes. The highly charged traditional SMA polymers are known to solubilize the cell membrane efficiently at a higher pH (pH 9) as compared to that at a neutral pH (pH 7) [28]. This solubilization effect has also been shown to be protein independent. Therefore, we wanted to examine how different pH conditions affect the efficacy of the pentyl-inulin to solubilize the *E. coli* membranes and, subsequently, determine the yield of FBD, Cyt-b5, and total *E. coli* membrane proteins. The solubilization was performed at three different pH values (7, 8, and 9), and the FBD protein bands were quantified from SDS-PAGE gel, and the Cyt-b5 was quantified by recording absorbance spectra (**Figs. 4** and **S3**). As shown in Figure 4A, the FBD protein band intensity showed no significant difference for pH 7 and pH 8. But, the observed FBD band intensity increased by ∼3.5 % upon increasing the pH to 9 (**Figs. 4A** and **S3**), suggesting a small improvement in the efficacy of the pentyl-inulin polymer in solubilizing the FBD-rich *E. coli* membranes. In the case of Cyt-b5, the yield of protein extraction increased by 15 % when the pH was increased from 7 to 9 (**Figs. 4(B, C)** and **S3**), indicating the effective membrane solubilization by the pentyl-inulin polymer at a higher pH. However, there was no difference in the amount of total membrane proteins solubilized at these three pH conditions (**Fig. 4D**). Overall, the results suggest that the buffer pH affect the pentyl-inulin-based solubilization of *E. coli* membranes in a protein-dependent manner and highlighting the suitability of non-ionic polymers for the solubilization of *E. coli* membranes to isolate a variety of bacterial membrane proteins at different pH conditions.

**Figure 4.**
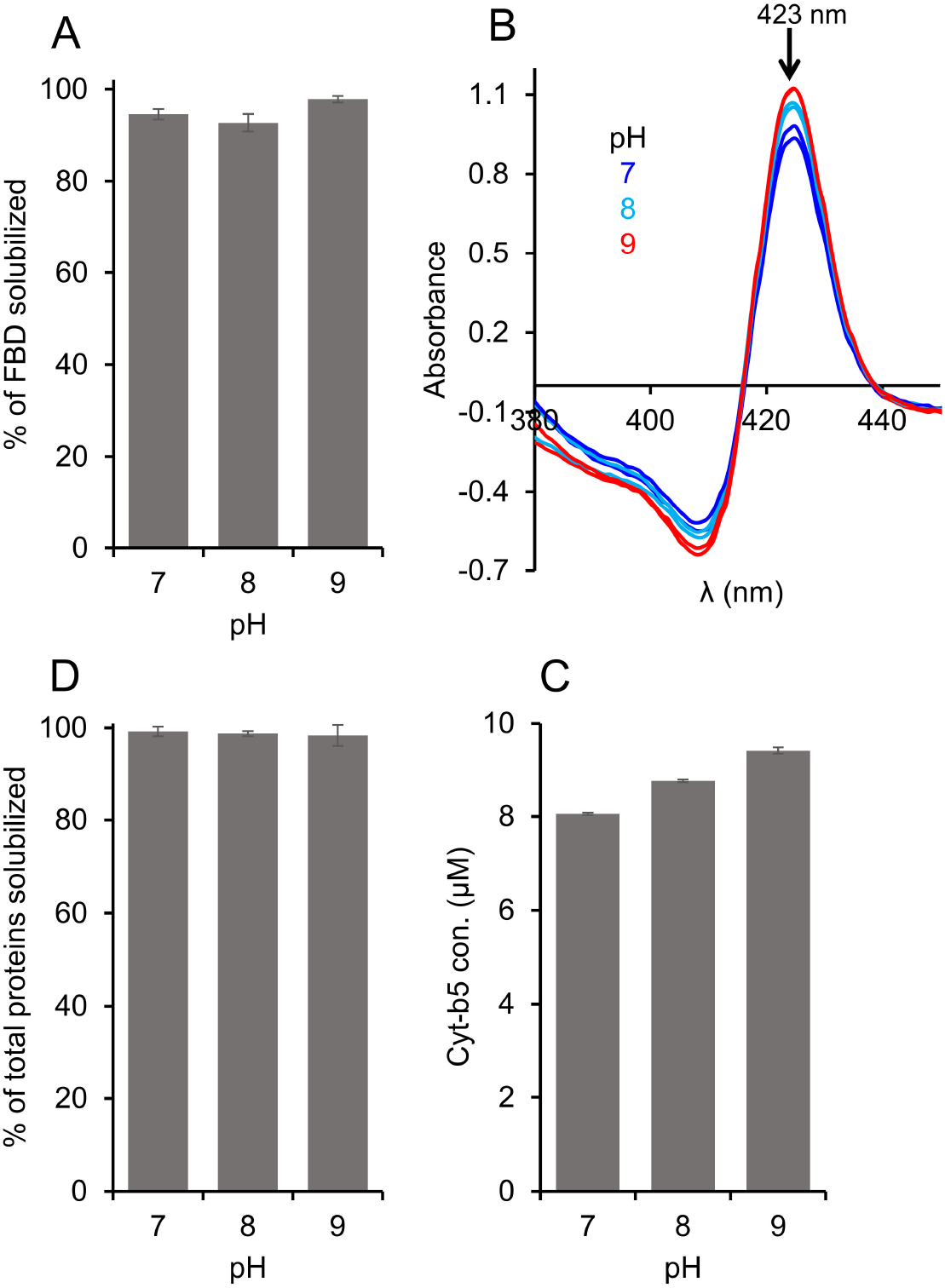
Effect of pH on the pentyl-inulin-based solubilization of *E. coli* membranes enriched with (**A**) FBD, (**B, C**) Cyt-b5, and (**D**) total *E. coli* membrane proteins. Cyt-b5 concentration was measured using absorbance spectra recorded at different pH conditions as indicated (B). The bar-graphs indicate the % of proteins in the polymer-solubilized samples estimated from protein band intensities in SDS-PAGE gels using ImageJ software. Error bars indicate the difference in the extent of solubilization between two independent experiments; data points are given as average values.

### 3.4. Ca^2+^ and Mg^2+^ ions affect the solubilization of E. coli membranes in a protein-dependent manner

SMA-based polymers are known to be unstable in the presence of divalent metal ions. Therefore, the negatively charged SMA polymers are not suitable for studying metal-binding biomolecules. The instability of SMA polymers is due to the presence of negatively charged carboxylic groups that strongly chelate with positively charged divalent metal ions. In contrast, the non-ionic pentyl-inulin contains no charged groups and is resistant to divalent metal ions [34]. Therefore, we wanted to examine the suitability of pentyl-inulin for the detergent-free isolation of membrane proteins in the presence of divalent metal ions.

The *E. coli* membranes enriched with FBD and Cyt-b5 were solubilized using 5 different concentrations of Mg^2+^ and Ca^2+^ ions (**Figs. 5** and **S4**). The intensity of the FBD protein band was not substantially changed when the solubilization was performed in the presence of Mg^2+^ up to 100 mM concentration (**Fig. 5A**). However, at higher Mg^2+^ concentrations (150 mM), the protein band intensity decreased by ∼16 %. In the presence of Ca^2+^, the reduction in the FBD intensity was more prominent. The intensity decreased by ∼18 % when Ca^2+^ level was increased up to 100 mM (**Fig. 5B**). Previous study indicates that pentyl-inulin:zwitterionic DMPC nanodiscs are stable in the presence of divalent metal ions up to 100 mM concentration [34]. The observed difference can be due to the difference in the lipid composition between synthetic DMPC nanodiscs [34] and *E. coli* native nanodiscs. When Ca^2+^ concentration was increased further to 150 mM, the FBD protein band intensity was decreased by 55 % (**Fig. 5B**), indicating decreased solubilization of membranes. This may be due to the excess of divalent ions in solution interacting with anionic lipids of FBD-enriched bacterial membranes, thus interfering with the integrity of the lipid bilayer surrounding FBD. The soluble domain of FBD itself is also highly anionic (−24 charge at pH 7.4), but whether its interaction with metal ions affects the overall yield of membrane solubilization is difficult to explain. In the case of Cyt-b5, the intensity of the protein band corresponding to Cyt-b5 increased when Mg^2+^ concentration was increased from 0 to 30 mM (**Fig. 5C**). Further, with the increase of [Mg^2+^] (up to 150 mM), there was no substantial change in the observed intensity of Cyt-b5 protein band (**Figs. 5C** and **S4**). This result suggests that the presence of Mg^2+^ (∼30 mM) improves the solubilization of Cyt-b5-enriched bacterial membranes. The influence of Ca^2+^ on the solubilization of membranes was somewhat similar to that of Mg^2+^ (**Figs. 5C, D**, and **S4**). The protein band intensity of Cyt-b5 increased to the maximum when the Ca^2+^ concentration was increased from 0 to 50 mM (**Fig. 5D**). But at higher Ca^2+^ concentrations (>100 mM), the intensity of several protein bands was decreased (**Fig. 5D** and **S4**). The change in the band intensities was more prominent in the Ca^2+^-containing samples when compared to the Mg^2+^-containing samples.

**Figure 5.**
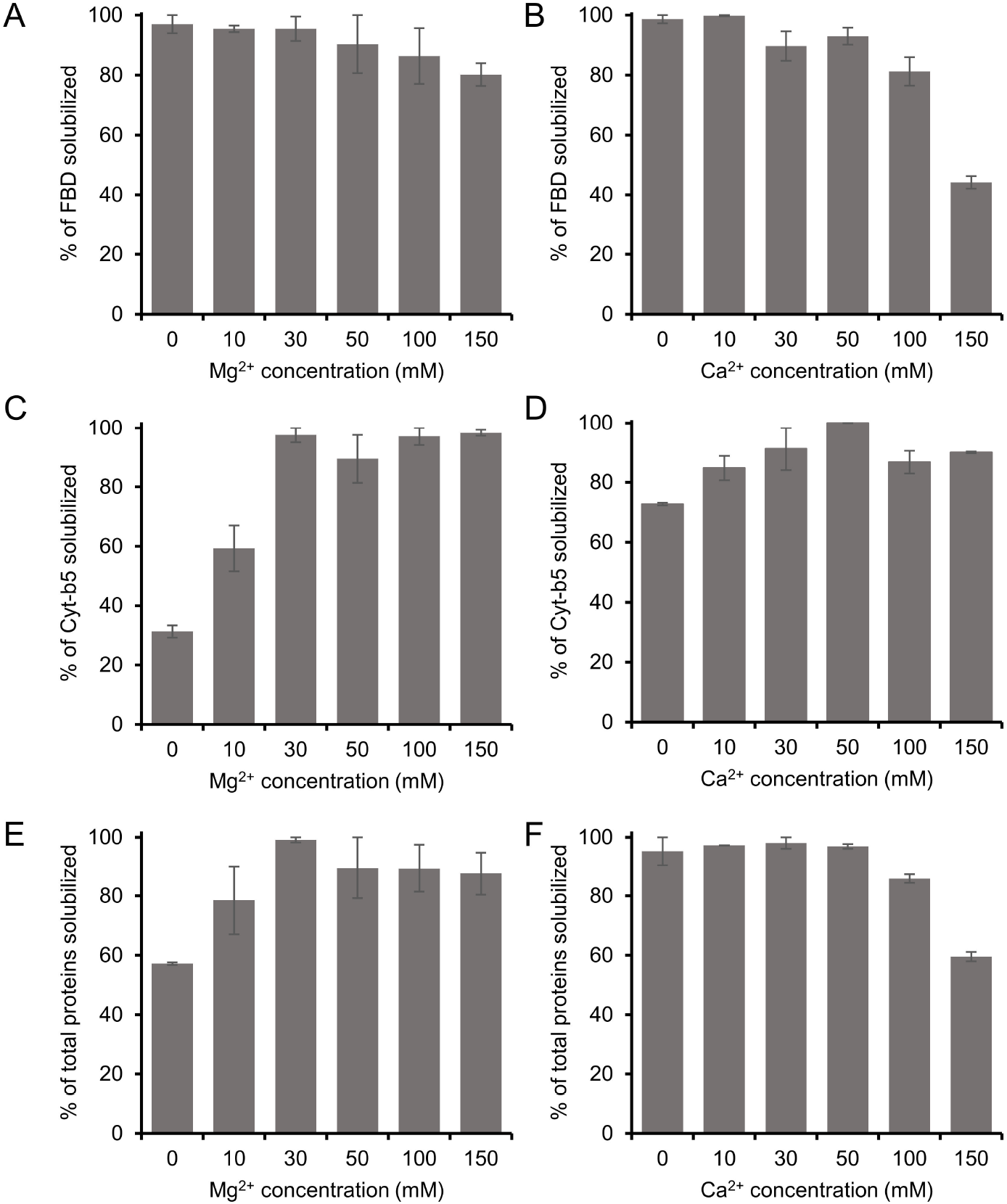
Influence of divalent metal ions (Mg^2+^ and Ca^2+^) on the solubilization of *E. coli* membranes enriched with (**A, B**) FBD, (**C, D**) Cyt-b5 and (**E, F**) total membrane proteins. The samples were prepared at a 1:1 w/w polymer-to-membrane ratio, and the solubilization was performed at 4 °C overnight in the presence of various concentrations of Mg^2+^ and Ca^2+^ ions, as indicated. The samples were analyzed by SDS-PAGE (Fig. S4) and quantified by using the Image Lab software. The protein band corresponding to Cyt-b5 is labelled, and the variations in the protein band intensity of *E. coli* membrane proteins at different metal ion concentrations are indicated with arrows. Only the well-resolved protein bands showing variation with different metal ion concentrations are indicated for clarity. The plots were generated from two independent experiments, and the error bars indicate the difference in the extent of solubilization between them. The data were normalized to the sample that showed the maximum solubilization.

In addition, the intensities of all solubilized membrane protein bands were quantified to evaluate the effect of metal ions on the membrane solubilization efficacy of the pentyl-inulin polymer. The total intensity of protein bands on the SDS-PAGE gel (corresponding to all membrane proteins) increased with the increase in the concentration of Mg^2+^ up to 30 mM (**Fig. 5E**). At higher Mg^2+^ concentrations (50 mM and above), there was only a small difference in the intensities when compared to lower concentrations of Mg^2+^ (**Fig. 5E**); this was due to a decrease in the intensity of some of *E. coli* cell membrane proteins (**Fig. S4**). The decrease in intensity or disappearance of some protein bands was possibly caused by the aggregation/precipitation of those membrane proteins associated with anionic lipids that are likely to interact with an excess of Mg^2+^ ions. On the other hand, there was no difference in the total intensity of protein bands up to 50 mM Ca^2+^ (**Fig. 5F** and **S4**). However, when the Ca^2+^ concentration increased above 50 mM, there was a substantial decrease in the total proteins’ band intensity, as seen on SDSPAGE gel (**Fig. S4**). In addition, some protein bands disappeared completely at higher Ca^2+^ concentrations (**Fig. S4**; indicated with arrows).

### 3.5. Effect of different functionalization of inulin polymer on the solubilization of E. coli membranes

FBD-enriched membranes were solubilized using inulin polymers functionalized with five different hydrophobic groups (butyl, pentyl, hexyl, benzyl and biphenyl) (**Fig. 1**). All of the samples were prepared using 1:1 (w/w) membrane-to-polymer ratios and the intensities of FBD protein bands on the SDS-PAGE gel were quantified (**Fig. S5**). We observed no statistical difference in the FBD protein band intensities from the samples solubilized using butyl or pentyl or hexyl or benzyl functionalized-inulins as analyzed from two independent solubilization experiments (**Fig. 6A**). But a ∼10 % increase was observed in the solubilization of total membrane proteins by inulin-hexyl compared to butyl-inulin (**Fig. 6B**). The biphenyl-inulin polymer also solubilized FBD-rich membranes (**Fig. S5**). However, due to its low-aqueous solubility, the data were not used to compare with that obtained for other inulin polymers. The commercial SMA and Sokalan-derived DIBMA polymers with a broad molecular weight distribution generally form smears on SDS-PAGE gels [42], thus limiting protein analysis. Interestingly, the inulin polymers functionalized with the aromatic benzyl or biphenyl groups with the degree of substitution of ∼0.3 – 0.5 did not affect the protein analysis by SDS-PAGE. This could be due to the free migration of polymer on the SDS-PAGE gel due to its low molecular weight (<4 kDa). Overall, the results reported in this study demonstrate that the inulin based polymers functionalized with different types of hydrophobic moieties are efficient in solubilizing the cell membranes and therefore suitable for detergent-free membrane protein isolation. However, it might be necessary to optimize the solubilization conditions to efficiently isolate any target membrane proteins using a specific polymer-type.

**Figure 6.**
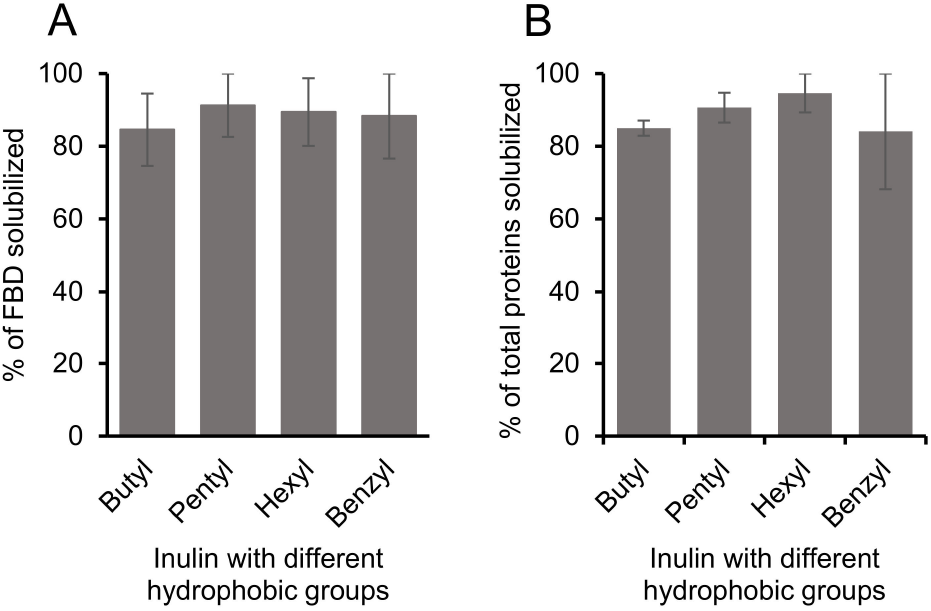
Influence of different hydrophobic groups on the solubilization of (**A**) FBD-rich membranes and (**B**) total membrane proteins. The plots were generated from two independent experiments, and the error bars indicate the difference in the extent of solubilization between them. The data were normalized to the sample that showed the maximum solubilization.

The polymers with high-charge density are not suitable for studies on membrane proteins whose charge is opposite to that of the polymer or for the protein-protein complexes that are stabilized by charge-charge interactions [17, 26]. Hence nanodiscs made with non-ionic polymers have been shown to be better suited to overcome such charge-based issues [29]. However, as discussed above, several other factors, including sample pH, temperature, polymer concentration and metal ions can affect the solubilization and nanodiscs formation efficacies of a given polymer. Therefore, it is useful to have knowledge on a given polymer’s membrane solubilizing ability in order to efficiently isolate the target membrane protein directly from the cell membrane. In addition to directly isolating target membrane proteins from *E. coli* and various cell types [10], the optimized membrane solubilization conditions can have a range of applications, such as the efficient isolation of low-expressing membrane proteins (GPCRs, cytochrome P450s) and amyloids from brain tissues. Additionally, one of the main applications of polymers is the effective preparation of native nanodiscs from various pathogenic and host cell membranes for treating various diseases. For example, Zhang and coworkers have isolated native nanodiscs from human red blood cell (RBC) membranes and showed their effective neutralization of hemolytic and cytotoxic effects caused by purified α-toxin or whole secreted proteins from methicillin-resistant *Staphylococcus aureus in vitro* [42]. In mice models, the native nanodiscs enriched with the RBC membranes conferred protection from *S. aureus* infection. Before performing such experiments, it was required to optimize the solubilization conditions (membrane-to-polymer ratio) to achieve an efficient nanodisc formation. The same research group has also isolated the membranes of *Pseudomonas aeruginosa* (the causative agent of pneumonia) as native nanodiscs using the SMA polymer to develop an effective antibacterial vaccine formulation [43]. The nanodiscs enriched with the outer membrane proteins (depending on the polymer) of *P. aeruginosa* elicited a strong immune response that protected a murine model from infection.

## 4. Conclusions

Although nanodiscs-forming synthetic polymers are increasingly used to directly isolate membrane proteins from the cell membranes and reconstitute them in native-membrane nanodiscs, the protocol and the conditions must be optimized for each system under investigation and can also vary with the type of polymer used. In this study, we have attempted to show how various solubilization conditions affect the solubilization efficacy of a non-ionic polymer, pentyl-inulin. The 1:1 (w/w) membrane:polymer ratio has the most efficient solubilization effect among the different polymer concentrations tested. Low temperatures (4 °C and 25 °C) showed better solubilization of membranes when compared to 37 °C. The use of sub-millimolar concentrations of divalent metal ions do not affect FBD/Cyt-b5-enriched membrane solubilization, signifying the use of a non-ionic polymer that does not directly interact with divalent metal ions. pH affected pentyl-inulin’s ability to solubilize different membrane proteins at different levels. In addition, inulin functionalized with different types of hydrophobic groups did not affect the solubilization of FBD-rich membranes. Inulin-based polymers do not affect the protein analysis/quantification by SDS-PAGE. Polymer’s solubilization efficacy can vary from one cell membrane type to another, which differs in the lipid composition and also could depend on the type of a target protein expressed. The efficacy of the polymer can be different when membrane proteins are expressed and purified from different types of bacteria or eukaryotic cells with different membrane lipid compositions. Therefore, the conditions examined in this study need not be optimal for the isolation of other membrane proteins. However, the approach and results reported in this study provide a broad idea about the different experimental and sample conditions that can be optimized to accomplish direct isolation of membrane proteins from the cell membrane and their functional reconstitution in polymer-based nanodiscs.

## CRediT authorship contribution statement

B.K. and A.R. designed the research. B.K. and G.S. optimized the approaches and carried out the experiments. T.R. synthesized and characterized the polymers. B.K. and G.S. processed, analyzed and curated the data. B.K., G.S. and A. R. interpreted the results. B.K. and A.R. wrote the manuscript. All authors have read and approved the final version of the manuscript. A. R. directed the project.

## Acknowledgment

We thank Joseph Marte for his help with the synthesis of one of the batches of pentyl-inulin polymer used in this study.

## References

[1] I.G. Denisov, S.G. Sligar, Nanodiscs for structural and functional studies of membrane proteins, Nat. Struct. Mol. Biol., 23 (23) 481–486.

[2] S.G. Sligar, I.G. Denisov, Nanodiscs: A toolkit for membrane protein science, Protein Sci., 30 (30) 297–315.

[3] I.G. Denisov, Y.V. Grinkova, A.A. Lazarides, S.G. Sligar, Directed self-assembly of monodisperse phospholipid bilayer nanodiscs with controlled size, J. Am. Chem. Soc., 126 (126) 3477–3487.

[4] U. Günsel, F. Hagn, Lipid nanodiscs for high-resolution NMR studies of membrane proteins, Chem. Rev., 122 (122) 9395–9421.

[5] F. Hagn, M. Etzkorn, T. Raschle, G. Wagner, Optimized phospholipid bilayer nanodiscs facilitate high-resolution structure determination of membrane proteins, J. Am. Chem. Soc., 135 (135) 1919–1925.

[6] M.L. Nasr, D. Baptista, M. Strauss, Z.J. Sun, S. Grigoriu, S. Huser, A. Plückthun, F. Hagn, T. Walz, J.M. Hogle, G. Wagner, Covalently circularized nanodiscs for studying membrane proteins and viral entry, Nat. Methods, 14 (14) 49–52.

[7] F. Hagn, M.L. Nasr, G. Wagner, Assembly of phospholipid nanodiscs of controlled size for structural studies of membrane proteins by NMR, Nat. Protoc., 13 (13) 79–98.

[8] S.C. Lee, T.J. Knowles, V.L. Postis, M. Jamshad, R.A. Parslow, Y.P. Lin, A. Goldman, P. Sridhar, M. Overduin, S.P. Muench, T.R. Dafforn, A method for detergent-free isolation of membrane proteins in their local lipid environment, Nat. Protoc., 11 (11) 1149–1162.

[9] J.M. Dörr, M.C. Koorengevel, M. Schäfer, A.V. Prokofyev, S. Scheidelaar, E.A.W. van der Cruijsen, T.R. Dafforn,M. Baldus, J.A. Killian, Detergent-free isolation, characterization, and functional reconstitution of a tetrameric K+ channel: The power of native nanodiscs, Proc. Natl. Acad. Sci. USA, 111 (111) 18607–18612.

[10] B. Krishnarjuna, A. Ramamoorthy, Detergent-free isolation of membrane proteins and strategies to study them in a near-native membrane environment, Biomolecules, 12 (12) 1076.

[11] C.J. Brown, C. Trieber, M. Overduin, Structural biology of endogenous membrane protein assemblies in native nanodiscs, Curr. Opin. Struct., 69 (69) 70–77.

[12] S. Lavington, A. Watts, Detergent-free solubilisation & purification of a G protein coupled receptor using a polymethacrylate polymer, BBA-Biomembranes, 1863 (1863) 183441.

[13] A.R. Long, C.C. O’Brien, K. Malhotra, C.T. Schwall, A.D. Albert, A. Watts, N.N. Alder, A detergent-free strategy for the reconstitution of active enzyme complexes from native biological membranes into nanoscale discs, BMC Biotechnol., 13 (13) 41.

[14] D.J.K. Swainsbury, S. Scheidelaar, R. van Grondelle, J.A. Killian, M.R. Jones, Bacterial reaction centers purified with styrene maleic acid copolymer retain native membrane functional properties and display enhanced stability, Angew. Chem. Int. Ed., 53 (53) 11803–11807.

[15] S. Gulati, M. Jamshad, Timothy J. Knowles, Kerrie A. Morrison, R. Downing, N. Cant, R. Collins, Jan B. Koenderink, Robert C. Ford, M. Overduin, Ian D. Kerr, Timothy R. Dafforn, Alice J. Rothnie, Detergent-free purification of ABC (ATP-binding-cassette) transporters, Biochem. J., 461 (461) 269–278.

[16] B. Krishnarjuna, T. Ravula, A. Ramamoorthy, Detergent-free isolation of CYP450-reductase’s FMN-binding domain in E.coli lipid-nanodiscs using a charge-free polymer, ChemComm, 58 (58) 4913–4916.

[17] B. Krishnarjuna, T. Ravula, A. Ramamoorthy, Detergent-free extraction, reconstitution and characterization of membrane-anchored cytochrome-b5 in native lipids, ChemComm, 56 (56) 6511–6514.

[18] A.O. Oluwole, B. Danielczak, A. Meister, J.O. Babalola, C. Vargas, S. Keller, Solubilization of membrane proteins into functional lipid-bilayer nanodiscs using a diisobutylene/maleic acid copolymer, Angew. Chem. Int. Ed., 56 (56) 1919–1924.

[19] B. Danielczak, M. Rasche, J. Lenz, E. Pérez Patallo, S. Weyrauch, F. Mahler, M.T. Agbadaola, A. Meister, J.O. Babalola, C. Vargas, C. Kolar, S. Keller, A bioinspired glycopolymer for capturing membrane proteins in native-like lipid-bilayer nanodiscs, Nanoscale, 14 (14) 1855–1867.

[20] M. Jamshad, J. Charlton, Y.-P. Lin, Sarah J. Routledge, Z. Bawa, Timothy J. Knowles, M. Overduin, N. Dekker, Tim R. Dafforn, Roslyn M. Bill, David R. Poyner, M. Wheatley, G-protein coupled receptor solubilization and purification for biophysical analysis and functional studies, in the total absence of detergent, Biosci. Rep., 35 (35) 1–10.

[21] M.C. Fiori, W. Zheng, E. Kamilar, G. Simiyu, G.A. Altenberg, H. Liang, Extraction and reconstitution of membrane proteins into lipid nanodiscs encased by zwitterionic styrene-maleic amide copolymers, Sci. Rep., 10 (10) 9940.

[22] D. Glueck, A. Grethen, M. Das, O.P. Mmeka, E.P. Patallo, A. Meister, R. Rajender, S. Kins, M. Räschle, J. Victor, C. Chu, M. Etzkorn, Z. Köck, F. Bernhard, J.O. Babalola, C. Vargas, S. Keller, Electroneutral polymer nanodiscs enable interference-free probing of membrane proteins in a lipid-bilayer environment, Small, 18 (18) 2202492.

[23] C. Sun, S. Benlekbir, P. Venkatakrishnan, Y. Wang, S. Hong, J. Hosler, E. Tajkhorshid, J.L. Rubinstein, R.B. Gennis, Structure of the alternative complex III in a supercomplex with cytochrome oxidase, Nature, 557 (557) 123–126.

[24] M. Parmar, S. Rawson, C.A. Scarff, A. Goldman, T.R. Dafforn, S.P. Muench, V.L.G. Postis, Using a SMALP platform to determine a sub-nm single particle cryo-EM membrane protein structure, BBA-Biomembranes, 1860 (1860) 378–383.

[25] D.J.K. Swainsbury, F.R. Hawkings, E.C. Martin, S. Musiał, J.H. Salisbury, P.J. Jackson, D.A. Farmer, M.P. Johnson, C.A. Siebert, A. Hitchcock, C.N. Hunter, Cryo-EM structure of the four-subunit Rhodobacter sphaeroides cytochrome bc_1_complex in styrene maleic acid nanodiscs, Proc. Natl. Acad. Sci. USA, 120 (120) e2217922120.

[26] T. Ravula, N.Z. Hardin, J. Bai, S.C. Im, L. Waskell, A. Ramamoorthy, Effect of polymer charge on functional reconstitution of membrane proteins in polymer nanodiscs, ChemComm, 54 (54) 9615–9618.

[27] A.H. Kopf, O. Lijding, B.O.W. Elenbaas, M.C. Koorengevel, J.M. Dobruchowska, C.A. van Walree, J.A. Killian, Synthesis and evaluation of a library of alternating amphipathic copolymers to solubilize and study membrane proteins, Biomacromolecules, 23 (23) 743–759.

[28] A.H. Kopf, J.M. Dörr, M.C. Koorengevel, F. Antoniciello, H. Jahn, J.A. Killian, Factors influencing the solubilization of membrane proteins from Escherichia coli membranes by styrene-maleic acid copolymers, BBA-Biomembranes, 1862 (1862) 183125.

[29] B. Krishnarjuna, S.C. Im, T. Ravula, J. Marte, R.J. Auchus, A. Ramamoorthy, Non-ionic inulin-based polymer nanodiscs enable functional reconstitution of a redox complex composed of oppositely charged CYP450 and CPR in a lipid bilayer membrane, Anal. Chem., 94 (94) 11908–11915.

[30] N.G. Brady, C.E. Workman, B. Cawthon, B.D. Bruce, B.K. Long, Protein extraction efficiency and selectivity of esterified styrene–maleic acid copolymers in thylakoid membranes, Biomacromolecules, 22 (22) 2544–2553.

[31] S. Scheidelaar, Martijn C. Koorengevel, Cornelius A. van Walree, Juan J. Dominguez, Jonas M. Dörr, J.A. Killian, Effect of polymer composition and pH on membrane solubilization by styrene-maleic acid copolymers, Biophys. J., 111 (111) 1974–1986.

[32] D.J.K. Swainsbury, S. Scheidelaar, N. Foster, R. van Grondelle, J.A. Killian, M.R. Jones, The effectiveness of styrene-maleic acid (SMA) copolymers for solubilisation of integral membrane proteins from SMA-accessible and SMA-resistant membranes, BBA-Biomembranes, 1859 (1859) 2133–2143.

[33] S.C.L. Hall, C. Tognoloni, G.J. Price, B. Klumperman, K.J. Edler, T.R. Dafforn, T. Arnold, Influence of poly(styrene-co-maleic acid) copolymer structure on the properties and self-assembly of SMALP nanodiscs, Biomacromolecules, 19 (19) 761–772.

[34] T. Ravula, A. Ramamoorthy, Synthesis, characterization, and nanodisc formation of non-ionic polymers, Angew. Chem. Int. Ed., 60 (60) 16885–16888.

[35] K.A. Morrison, A. Akram, A. Mathews, Z.A. Khan, J.H. Patel, C. Zhou, D.J. Hardy, C. Moore-Kelly, R. Patel, V. Odiba, T.J. Knowles, M.-u.-H. Javed, N.P. Chmel, T.R. Dafforn, A.J. Rothnie, Membrane protein extraction and purification using styrene–maleic acid (SMA) copolymer: effect of variations in polymer structure, Biochem. J., 473 (473) 4349–4360.

[36] B. Danielczak, A. Meister, S. Keller, Influence of Mg2+ and Ca2+ on nanodisc formation by diisobutylene/maleic acid (DIBMA) copolymer, Chem. Phys. Lipids, 221 (221) 30–38.

[37] I.G. Denisov, Y.V. Grinkova, M.A. McLean, T. Camp, S.G. Sligar, Midazolam as a probe for heterotropic drug-drug interactions mediated by CYP3A4, Biomolecules, 12 (2022).

[38] I.G. Denisov, Y.V. Grinkova, J.L. Baylon, E. Tajkhorshid, S.G. Sligar, Mechanism of drugdrug interactions mediated by human cytochrome P450 CYP3A4 monomer, Biochemistry, 54 (54) 2227–2239.

[39] C. Barnaba, K. Gentry, N. Sumangala, A. Ramamoorthy, The catalytic function of cytochrome P450 is entwined with its membrane-bound nature, F1000Res, 6 (6) 662.

[40] S.B. Mulrooney, L. Waskell, High-level expression in Escherichia coli and purification of the membrane-bound form of cytochrome b_5_, Protein Expr. Purif, 19 (19) 173–178.

[41] B. Krishnarjuna, J. Marte, T. Ravula, A. Ramamoorthy, Enhancing the stability and homogeneity of non-ionic polymer nanodiscs by tuning electrostatic interactions, J. Colloid Interface Sci., 634 (634) 887–896.

[42] L. Sun, D. Wang, I. Noh, R.H. Fang, W. Gao, L. Zhang, Synthesis of erythrocyte nanodiscs for bacterial toxin neutralization, Angewandte Chemie (International ed. in English), (2023) e202301566.

[43] I. Noh, Z. Guo, J. Zhou, W. Gao, R.H. Fang, L. Zhang, Cellular nanodiscs made from bacterial outer membrane as a platform for antibacterial vaccination, ACS Nano, 17 (17) 1120–1127.

