## Supporting Information for "Factors influencing the detergent-free membrane protein isolation using synthetic nanodisc-forming polymers"

[@]**Corresponding Author**

Ayyalusamy Ramamoorthy

*equal contribution

### Amino acid sequences of FBD and Cyt-b5

*Rat flavin mononucleotide binding domain (FBD) of cytochrome P450 reductase*

```
MGDSHEDTSATMPEAVAEEVSLFSTTDMVLFSLIVGVLTYWFIFRKKKEEIPEFSKIQTTAPPV
KESSFVEKMKKTGRNIIVFYGSQTGTAEEFANRLSKDAHRYGMRGMSADPEEYDLADLSSLPEI
DKSLVVFCMATYGEGDPTDNAQDFYDWLQETDVDLTGVKFAVFGLGNKTYEHFNAMGKYVDQRL
EQLGAQRIFELGLGDDDGNLEEDFITWREQFWPAVCEFFGVEATGEEHHHHHH
```

*Rabbit cytochrome-b5*

```
GHHHHHHAAQSDKDVKYYTLEEIKKHNHSKSTWLILHHKVYDLTKFLEEHPGGEEVLREQAGGD
ATENFEDVGHSTDARELSKTFIIGELHPDDRSKLSKPMETLITTVDSNSSWWTNWVIPAISALI
VALMYRLYMADD
```

The transmembrane domains are underlined.

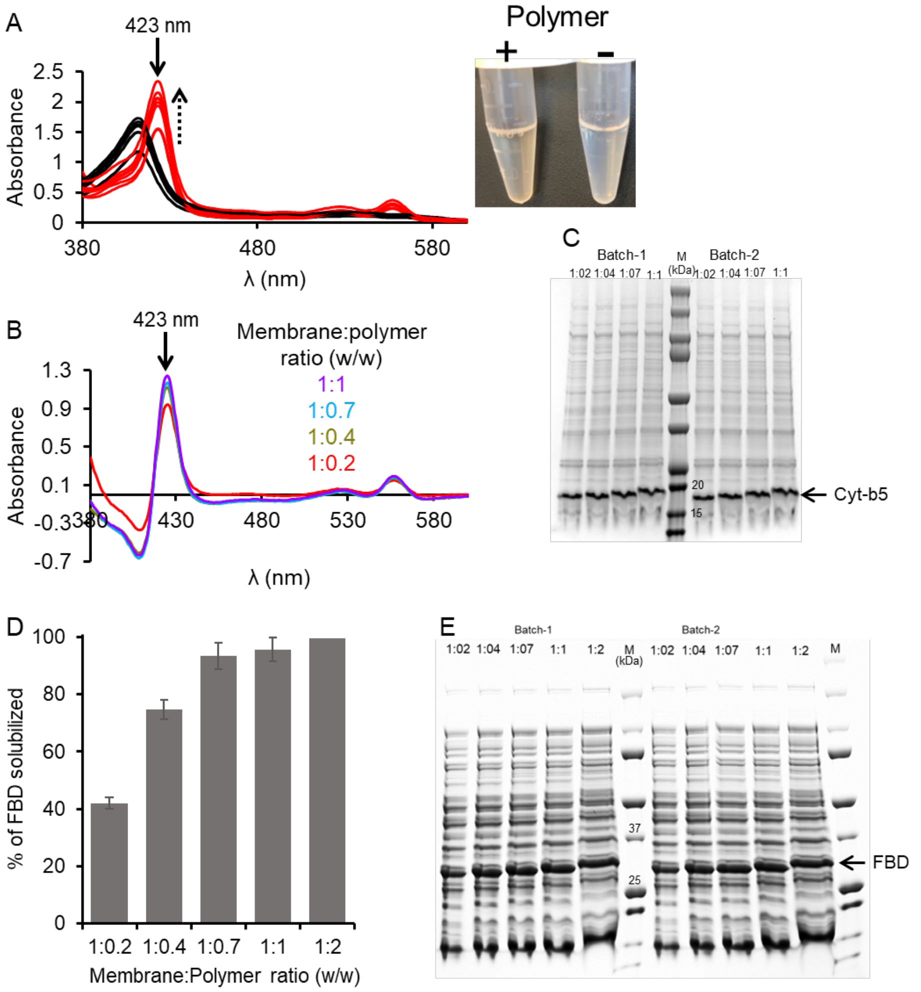


**Figure S1.** (**A**) Absorbance spectra of oxidized (black) and sodium dithionate reduced Cyt-b5 in *E. coli* lipid-pentyl-inulin nanodiscs. The samples were made using 4 different membrane:polymer (w/w) ratios as indicated. The absorbance peak intensity change at 423 nm is indicated with an upside dotted arrow. *Picture*: The light-red colored (left-side) and colorless (right-side) solutions in tubes are supernatants after centrifugation of Cyt-b5-enriched cell membranes with and without the polymer, respectively. (**B**) Difference absorbance spectra (reduced minus oxidized) of pentyl-inulin-solubilized *E.coli* membranes enriched with a ~15.7-kDa rabbit cytochrome-b5 showing the maximal absorbance differences at 409, 423, 526 and 556 nm. The data were collected on the solubilization samples prepared using 4 different polymer concentrations as indicated. (**C**) SDS-PAGE analysis of pentyl-inulin-solubilized Cyt-b5-rich *E. coli* cell membranes. (**D**) Bar plot depicting the percentage of 27.8-kDa FBD protein band intensities at 5 different membrane:polymer ratios (w/w) as indicated. (**E**) SDS-PAGE analysis of pentyl-inulin-solubilized FBD-rich *E. coli* cell membranes. Both Cyt-b5 and FBD are anchored to the lipid membrane via a transmembrane helical domain. The solubilization experiments were performed in duplicates (Batches 1 and 2). Cyt-b5 and FBD proteins have been directly isolated using SMA-EA and pentyl-inulin polymers, respectively [1-4].

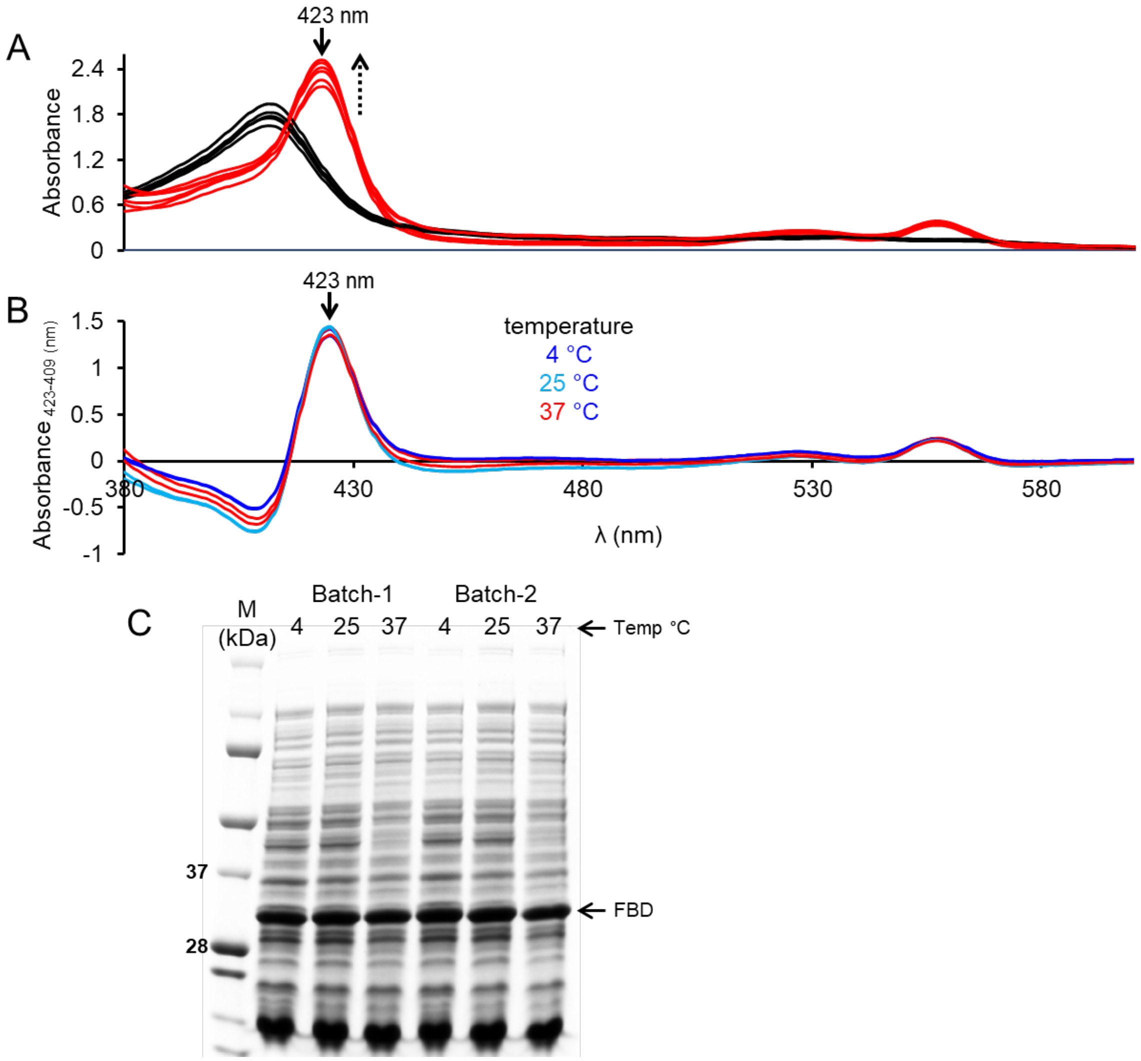


**Figure S2. (A)** Absorbance spectra of oxidized (black) and sodium dithionate reduced Cyt-b5 in *E. coli* lipid-pentyl-inulin nanodiscs. The samples were made using 1:1 membrane:polymer (w/w) ratio and solubilization was performed at 3 different temperatures as indicated. The absorbance peak intensity change at 423 nm is indicated with upside dotted arrow. (**B**) Difference absorbance spectra (reduced minus oxidized) of pentyl-inulin-solubilized *E.coli* membranes enriched with a ~15.7-kDa rabbit cytochrome-b5 showing the maximal absorbance differences at 409 and 423 nm. The data were collected on the solubilization samples prepared at 3 different temperature conditions as indicated. (**C**) SDS-PAGE analysis of pentyl-inulin-solubilized 27.8-kDa FBD-rich *E. coli* cell membranes obtained at different temperature conditions. M denotes the protein marker. The solubilization experiments were performed in duplicates (Batches 1 and 2).

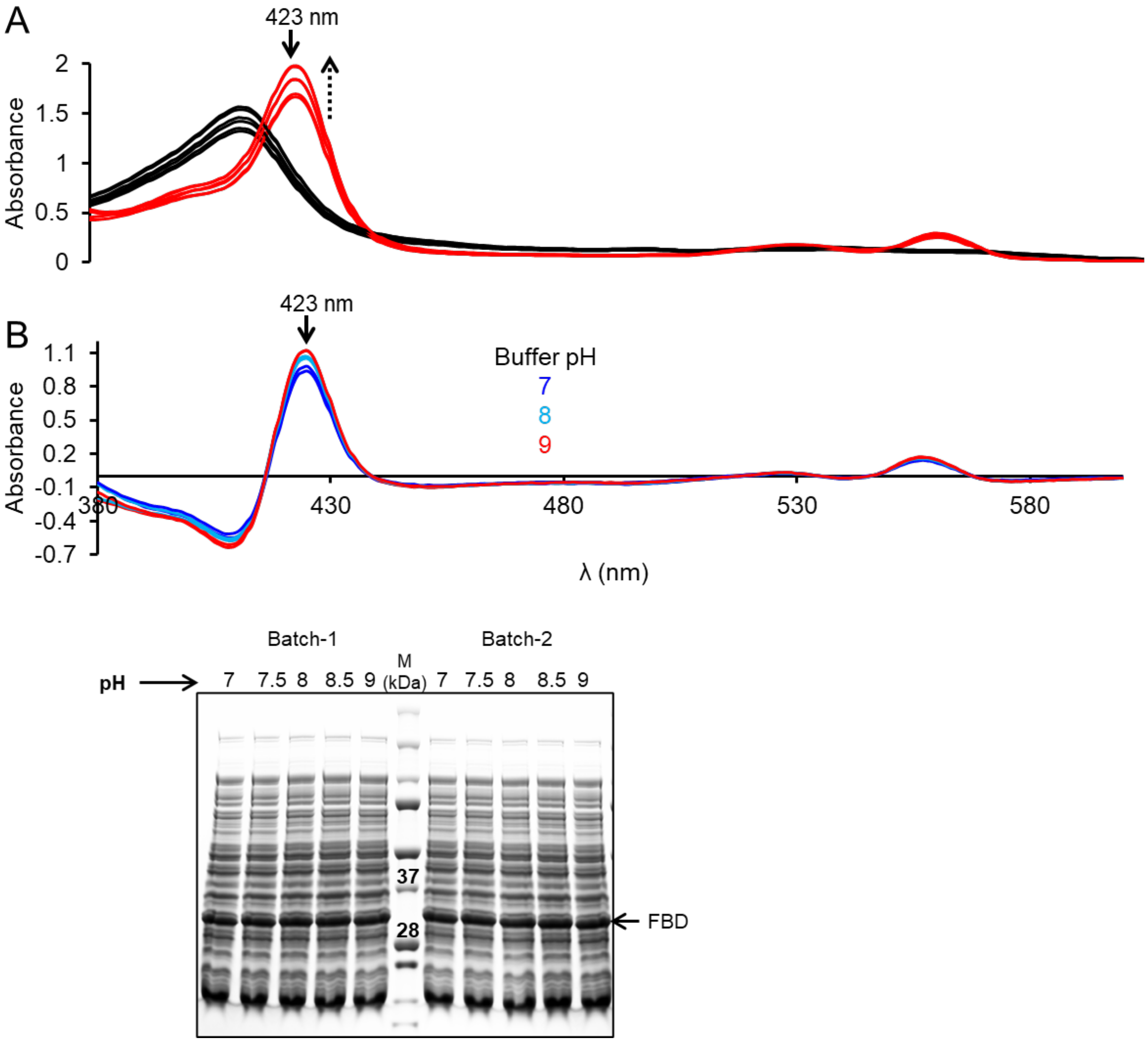


**Figure S3. (A)** Absorbance spectra of oxidized (black) and sodium dithionate reduced Cyt-b5 in *E. coli* lipid-pentyl-inulin nanodiscs. The samples were made using 1:1 membrane:polymer (w/w) ratio and solubilization was performed at 3 different pH conditions as indicated. The absorbance peak intensity change at 423 nm is indicated with an upside dotted arrow. (**B**) Difference absorbance spectra (reduced minus oxidized) of pentyl-inulin-solubilized *E.coli* membranes enriched with a ~15.7-kDa rabbit cytochrome-b5 showing the maximal absorbance differences at 409 and 423 nm. The data were collected on the solubilization samples prepared at 3 different pH conditions as indicated. (**C**) SDS-PAGE analysis of pentyl-inulin-solubilized 27.8-kDa FBD-rich *E. coli* cell membranes at different pH conditions. M denoted the protein marker. The solubilization experiments were performed in duplicates (Batches 1 and 2).

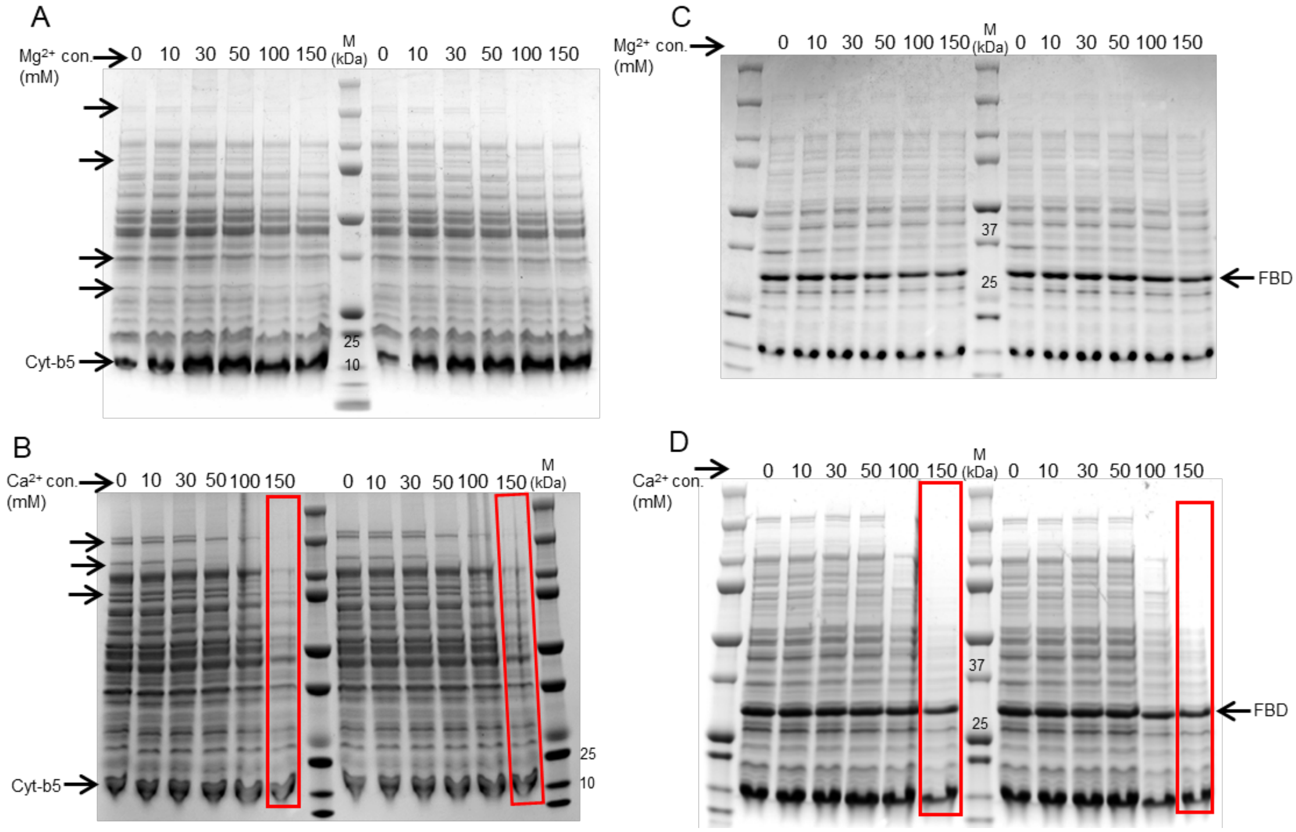


**Figure S4.** SDS-PAGE analysis of Cyt-b5 (**A, B**) and FBD-enriched (**C, D**) *E. coli* membranes that were solubilized using pentyl-inulin at the indicated concentrations of divalent metal ions. A and B are Cyt-b5 samples in the presence of $MgCl_2$ and $CaCl_2$, respectively. C and D are FBD samples in the presence of $MgCl_2$ and $CaCl_2$, respectively. The protein bands corresponding to Cyt-b5 and FBD are labelled. The metal ion concentrations are indicated. The lanes highlighted in (B) and (D) with red boxes indicate a significant decrease in the intensity of protein bands at higher concentrations of divalent metal ions. The solubilization experiments were performed in duplicates.

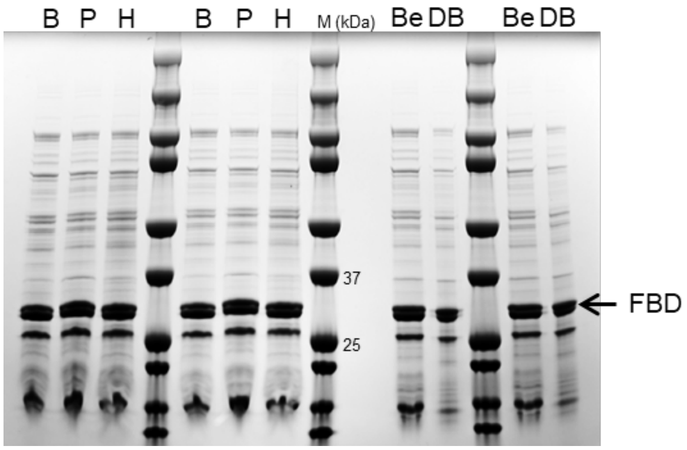


**Figure S5.** SDS-PAGE analysis of FBD-enriched *E. coli* membranes that were solubilized using inulin based polymers functionalized with butyl (B), pentyl (P), hexyl (H), benzyl (Be) or di-benzyl (DB) hydrophobic groups. The solubilization experiments were performed in duplicates.

### References


[1] B. Krishnarjuna, A. Ramamoorthy, Detergent-free isolation of membrane proteins and strategies to study them in a near-native membrane environment, Biomolecules, 12 (2022) 1076.
[2] B. Krishnarjuna, T. Ravula, A. Ramamoorthy, Detergent-free extraction, reconstitution and characterization of membrane-anchored cytochrome-b5 in native lipids, ChemComm, 56 (2020) 6511-6514.
[3] T. Ravula, A. Ramamoorthy, Synthesis, characterization, and nanodisc formation of non-ionic polymers, Angew. Chem. Int. Ed., 60 (2021) 16885-16888.
[4] T. Ravula, S.K. Ramadugu, G. Di Mauro, A. Ramamoorthy, Bioinspired, size-tunable self-assembly of polymer-lipid bilayer nanodiscs, Angew. Chem. Int. Ed., 56 (2017) 11466-11470.
